# piR-bmo-796514 Facilitates the Proliferation of Exogenous DNA Virus (Baculovirus) by Targeting the Host E3 Ubiquitin Ligase RNF181

**DOI:** 10.1101/2025.06.08.658493

**Authors:** Junming Xia, Shigang Fei, Wenjie Luo, Minyang Zhou, Yibing Kong, Yigui Huang, Luc Swevers, Min Feng

## Abstract

PIWI-interacting RNAs (piRNAs), a class of 23-31 nucleotide non-coding RNAs, are known for silencing transposons and endogenous retroviruses that reside in animal genomes. However, the mechanisms by which host piRNAs affect exogenous viral infections, particularly those by DNA viruses, remain poorly understood. Here, we demonstrate that infection by Bombyx mori nucleopolyhedrovirus (BmNPV), a large DNA virus, induces significant upregulation of silkworm host piR-bmo-796514, which facilitates viral proliferation by suppressing the expression of E3 ubiquitin ligase RNF181. We further reveal that RNF181 exerts antiviral activity through ubiquitin-mediated degradation of Integrin α2b-like, a cellular membrane protein that interacts with viral GP64 protein to mediate BmNPV entry. This study unveils a previously unrecognized regulatory axis connecting host derived piRNAs with exogenous DNA virus infection, providing further mechanistic insights into the modulation of exogenous viral pathogenesis through the reprogramming of the piRNA pathway. Our findings not only advance the understanding of the immune escape mechanism of exogenous viruses but also provide new insights for the development of oligonucleotide antiviral drugs that target proviral piRNAs.

## Introduction

Piwi-interacting RNAs (piRNAs) represent a distinct class of small non-coding RNAs that are fundamentally different from small interfering RNAs (siRNAs) and microRNAs (miRNAs) in terms of their biogenesis, molecular characteristics, and functional roles in cellular processes^1^. Typically ranging from 24 to 31 nucleotides in length, piRNAs are distinguished by the presence of 2′-O-methyl modifications at their 3′ termini, a feature that enhances their stability and functional specificity^2^. Unlike siRNAs and miRNAs, piRNAs are generated from precursor transcripts originating from intergenic regions known as piRNA clusters through a Dicer-independent pathway^3^. These clusters are densely populated with transposable elements (TEs), underscoring the primary role of piRNAs in silencing TE activity and maintaining genomic integrity, particularly during gametogenesis, where they are essential for preserving fertility^4^. The remarkable diversity of piRNA sequences, stemming from their varied processing mechanisms and origins within TE-rich genomic loci, sets them apart from other classes of non-coding RNAs^4,5^. This diversity not only reflects their evolutionary adaptation to counteract a wide array of TEs but also positions piRNAs as the most abundant and functionally versatile category of non-coding RNAs in the genome^6^.

In recent years, emerging evidence has expanded the functional repertoire of piRNAs beyond their canonical roles, implicating them in diverse biological processes, including cancer progression, epigenetic regulation, and host-pathogen interactions^6,7^. Notably, the involvement of piRNAs in antiviral defense has garnered increasing attention, particularly in the context of RNA viruses^8,9^. The first indications that the piRNA pathway might play a role in suppressing exogenous viruses came from research on mosquitoes, providing evidence for the potential importance of this pathway in the antiviral defense mechanisms of mosquitoes^10–13^.

Despite these advances, significant gaps remain in our understanding of piRNA-mediated antiviral mechanisms. First, most studies have focused on RNA viruses, leaving the role of piRNAs during exogenous DNA virus infections largely unexplored. Given the distinct replication strategies and host interactions of DNA viruses, it is unclear whether piRNAs exert similar or distinct regulatory functions in this context. Second, while some host derived piRNAs have been shown to directly target viral sequences, the potential indirect regulatory roles of piRNAs, such as modulating host factors critical for viral replication-remain poorly understood^14,15^. Third, the molecular mechanisms by which viral infections alter host piRNA expression profiles, and how these changes impact viral pathogenesis, are still enigmatic. Addressing these questions is critical for elucidating the complex interplay between host piRNA pathways and exogenous viral infections.

In addition to the well-established models of *Drosophila melanogaster* and aedine mosquitoes, significant progress in piRNA research has been achieved in other insect systems, particularly the silkworm. Especially important was the identification of piRNAs as key determinants of sex determination, providing novel insights into the role of piRNAs in sex chromosome function and reproductive development^16^. Silkworm cell lines, particularly the BmN4 cell line derived from silkworm embryos, have been instrumental in elucidating the molecular mechanisms underlying piRNA biogenesis and function^17,18^. Notably, the ping-pong amplification loop, a conserved mechanism for piRNA amplification, has been extensively studied in BmN4 cells, where it is mediated by the silkworm PIWI proteins BmAgo3 and Siwi^19,20^.

The major pathogen that causes disease in silkworms is the baculovirus, Bombyx mori nucleopolyhedrovirus (BmNPV). BmNPV and other baculoviruses are characterized by large (80-180 kb) circular dsDNA genomes encoding more than 100 genes that become expressed during different phases of a complex life cycle^21^. Intriguingly, our previous research found that after BmNPV infection in silkworms, no virus-derived piRNAs were detected in the fat body and midgut, but the expression of a large number of host derived piRNAs was significantly altered^22^. Moreover, based on silencing and over-expression of Siwi and BmAgo3 in BmN cells and silkworm larva, it was confirmed that the silkworm piRNA pathway could promote BmNPV proliferation^23^, in contrast to the proposed antiviral function of piRNAs against RNA viruses in mosquitoes and lepidopteran cells^18,24^. However, the specific mechanism by which the piRNA pathway in silkworm promotes BmNPV proliferation is still unclear.

The proviral role of PIWI-class proteins raises interest since a re-organization of the nucleus takes place during baculovirus infection^25^, which could affect the expression of piRNA clusters in the silkworm genome^22,26^. Most apparent is a nuclear viral substructure called the virogenic stroma that functions as a viral factory for viral DNA replication and transcription as well as nucleocapsid assembly^27^. In parallel, the cellular chromatin becomes condensed and marginalized to the nuclear periphery, also known as virus-induced reorganization of cellular chromatin^28,29^. A recent study indicates that BmNPV usurps the transcriptional regulatory machinery at the super-enhancers in the host cells’ chromatin to inhibit antiviral gene expression and promote the organization of the BmNPV genomes in the nuclei (Zhao et al., 2025). While marginalization to the nuclear periphery is expected to result in repression of cellular transcription, it is well known that many exceptions exist and that the virus is able to up-regulate certain categories of genes involved with metabolism and stress response at 6 and 12 hpi^30^. Of note is the differential expression of transposons that is stimulated by baculovirus infection^31,32^ and integration of transposons into baculoviral genomes is considered a potential factor for horizontal transmission of DNA between insects^33^. The differential silencing/expression of transposons during infection with BmNPV and other baculoviruses may be caused by the modulation of the piRNA pathway by BmNPV and other baculoviruses.

The separation of baculovirus genomes from the cellular chromatin to different nuclear compartments is thought to reflect the different conditions required for viral DNA replication, i.e. in the absence of deposition of histones^29^. In addition, it has been speculated that the re-organization of the cellular chromatin corresponds to a process of de-differentiation and may require the assembly of virus-modified chromatin remodeling complexes^34^. Recently, the role of the piRNA pathway and PIWI-class proteins in the regulation of cellular processes such as stem cell maintenance, genome re-organization, inflammation, ageing and cancer has become increasingly apparent^35–38^. In mammals, piRNAs have become important biomarkers for diseases, including viral infections, underscoring the need to unravel the mechanisms by which they play a role in pathogenesis^39,40^. The unraveling of piRNA and PIWI-class protein function during BmNPV infection will therefore contribute to the better understanding of the reprogramming of cellular processes by viruses.

In this study, utilizing the silkworm as a model organism, we have focused on a host derived piRNA, piR-bmo-796514, which is significantly induced by BmNPV infection and can promote (rather than inhibit) the proliferation of exogenous DNA viruses. Importantly, our findings demonstrated that piR-bmo-796514 promoted BmNPV proliferation by suppressing the expression of the E3 ubiquitin ligase RNF181, a homolog of human RNF181 (Ring finger 181. Brophy et al., 2008). Concurrently, we discovered that RNF181 exerted its antiviral function against BmNPV through the ubiquitination and subsequent degradation of the transmembrane receptor Integrin α2b-like, which interacts with the BmNPV GP64 envelope protein to facilitate viral entry. Collectively, this study provides a new perspective on the immune escape of exogenous viruses mediated by specific piRNA targeting host factors.

## Results

### piR-bmo-796514 facilitates BmNPV proliferation

In a previous study, we documented the response of host derived piRNAs piRNAs in the fat body and midgut of the silkworm to BmNPV infection^22^. Remarkably, piR-bmo-796514 exhibited a significant upregulation in the silkworm fat body after BmNPV infection (Fig. 1a). Subsequently, we monitored the expression of piR-bmo-796514 in the fat body and BmN cells at multiple time points following BmNPV infection. Our findings revealed a significant upregulation of piR-bmo-796514 in both the fat body and BmN cells at 24 hpi but not at 48 hpi (Fig. 1b, c). These results suggested that piR-bmo-796514 was involved in the process of BmNPV infection in the host cells and might play a role in the host responses.

**Fig. 1:**
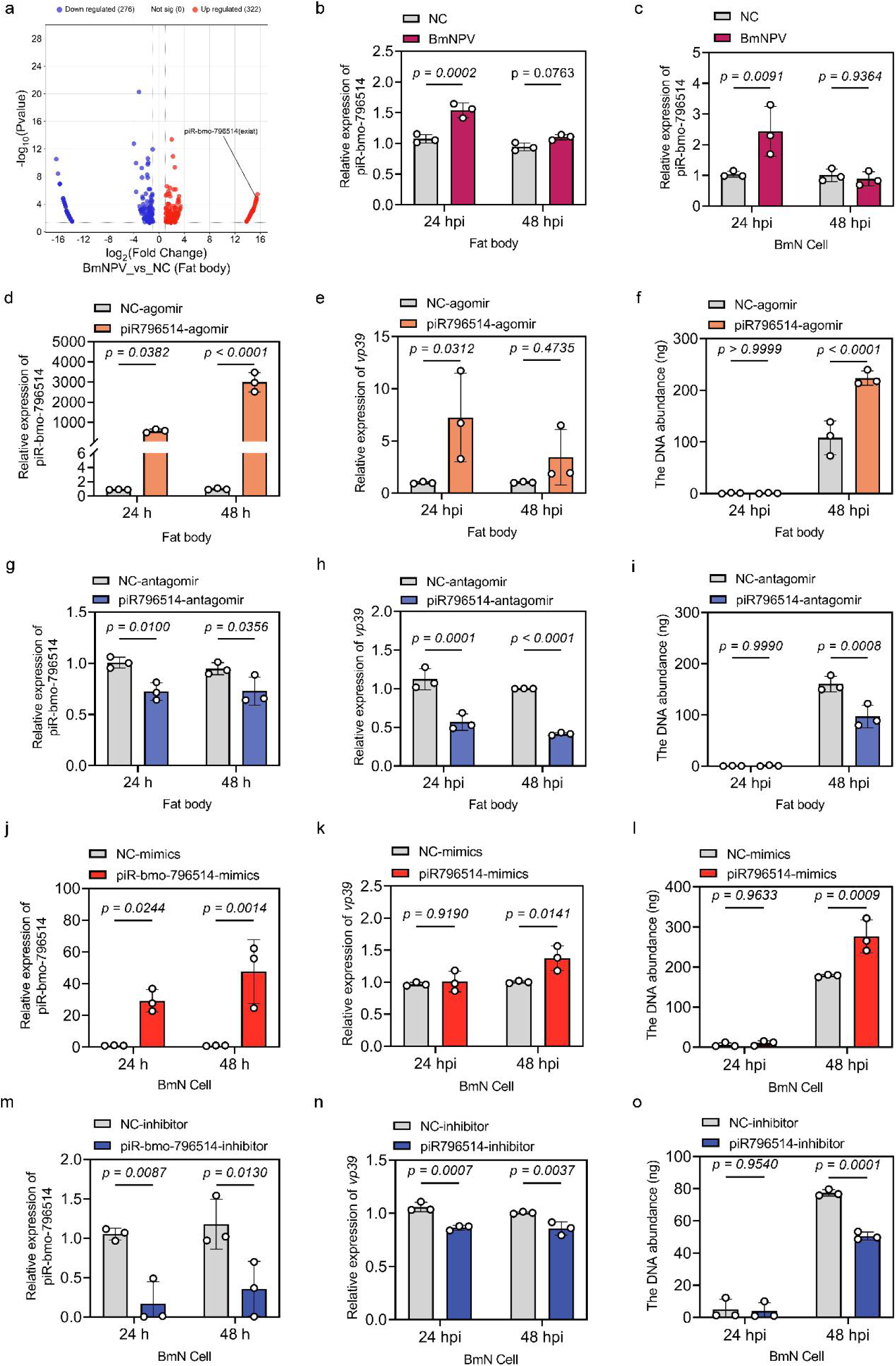
The piR-bmo-796514 is induced by BmNPV infection and promotes BmNPV proliferation. (a) Detection of differentially expressed (DE) piRNAs in the fat body of silkworms following BmNPV infection. (b, c) Quantitative detection of piR-bmo-796514 expression in the fat body of silkworms and BmN cells after BmNPV infection. (d) Detection of piR-bmo-796514 expression in the fat body of silkworm larvae after injection with piR-bmo-796514-agomir. (e) Detection of the transcriptional level of the viral *vp39* gene in the fat body of silkworm larvae after injection with piR-bmo-796514-agomir and subsequent BmNPV infection. (f) Detection of the DNA load of the viral *gp41* gene in the fat body of silkworm larvae after injection with piR-bmo-796514-agomir and subsequent BmNPV infection. (g) Detection of piR-bmo-796514 expression in the fat body of silkworm larvae after injection with piR-bmo-796514-antagomir. (h) Detection of the transcriptional level of the viral *vp39* gene in the fat body of silkworm larvae after injection with piR-bmo-796514-antagomir and subsequent BmNPV infection. (i) Detection of the DNA load of the viral *gp41* gene in the fat body of silkworm larvae after injection with piR-bmo-796514-antagomir and subsequent BmNPV infection. (j) Detection of piR-bmo-796514 expression in BmN cells after transfection with piR-bmo-796514-mimics. (k) Detection of the transcriptional level of the viral *vp39* gene in BmN cells after transfection with piR-bmo-796514-mimics and subsequent BmNPV infection. (l) Detection of the DNA load of the viral *gp41* gene in BmN cells after transfection with piR-bmo-796514-mimics and subsequent BmNPV infection. (m) Detection of piR-bmo-796514 expression in BmN cells after transfection with piR-bmo-796514-inhibitor. (n) Detection of the transcriptional level of the viral *vp39* gene in BmN cells after transfection with piR-bmo-796514-inhibitor and subsequent BmNPV infection. (o) Detection of the DNA load of the viral *gp41* gene in BmN cells after transfection with piR-bmo-796514-inhibitor and subsequent BmNPV infection. *p*-value <0.05 was considered as statistically significant.

A analyzing the genomic locus corresponding to piR-bmo-796514 revealed that piR-bmo-796514 did not belong to any annotated piRNA cluster (Supplementary Fig. 1). Alignment of the piR-bmo-796514 sequence against the silkworm genome identified five specific potential loci in the silkworm genome, four of which have not yet been annotated, and another gene (BMSK0002231) (Supplementary Fig. 1a, b) which is not responsive to BmNPV infection in fat body (Supplementary Fig. 1c) and BmN cells (Supplementary Fig. 1d). Further analysis of the piRBase database showed that piR-bmo-796514 binds to BmAgo3 in *Bombyx mori* BmN4 cells (Supplementary Fig. 2a). Our RNA immunoprecipitation (RIP) assays in BmN cells also confirmed the specific association of piR-bmo-796514 with BmAgo3 (Supplementary Fig. 2b, c).

To further explore the influence of piR-bmo-796514 on the proliferation of BmNPV, fifth-instar silkworm larvae were injected with piR-bmo-796514-agomir and subsequent analysis of fat body samples at 24 and 48 hours post-injection revealed a significant upregulation of piR-bmo-796514 expression (Fig. 1d). Furthermore, when piR-bmo-796514-agomir was co-injected with the BmNPV, the transcriptional level of the viral *vp39* gene was significantly increased at 24 hpi (Fig. 1e), and the DNA load of the BmNPV *gp41* gene was significantly increased at 48 hpi (Fig. 1f). These results strongly suggested that the overexpression of piR-bmo-796514 within the fat body of silkworm larvae exerted a promotional effect on BmNPV proliferation. Conversely, to assess the impact of reduced piR-bmo-796514 expression, we injected piR-bmo-796514-antagomir into the silkworm larvae. Analysis of fat body samples after injection of piR-bmo-796514-antagomir at 24 and 48 h demonstrated a substantial downregulation of piR-bmo-796514 expression (Fig. 1g). Co-injection of the virus with piR-bmo-796514-antagomir led to a significant decline in the transcriptional level of the BmNPV *vp39* gene at 24 and 48 hpi (Fig. 1h). Furthermore, at 48 hpi, there was a significant reduction in the DNA load of the BmNPV *gp41* gene (Fig. 1i). These findings implied that the suppression of piR-bmo-796514 in the fat body of silkworm larvae could effectively inhibit BmNPV proliferation.

We further evaluated the influence of piR-bmo-796514 on the proliferation of BmNPV in BmN cells. After transfection with piR-bmo-796514-mimics in BmN cells, a substantial increase in the expression level of piR-bmo-796514 was detected at 24 and 48 h post transfection (Fig. 1j). Subsequently, 24 hours post-transfection with piR-bmo-796514-mimics, BmN cells were infected with BmNPV. The results revealed a significant upregulation in the transcriptional level of the BmNPV *vp39* gene at 48 hpi (Fig. 1k). Meanwhile, the DNA load of the BmNPV *gp41* gene was also significantly elevated at 48 hpi (Fig. 1l). These findings strongly suggested that the overexpression of piR-bmo-796514 could facilitate the proliferation of BmNPV in BmN cells.

Conversely, to investigate the impact of reduced piR-bmo-796514 expression, BmN cells were transfected with a piR-bmo-796514 inhibitor, resulting in a decrease in piR-bmo-796514 expression (Fig. 1m). After infecting the transfected cells with BmNPV at 24 hours post-transfection, we observed a significant downregulation in the transcriptional level of the viral *vp39* gene at both 24 and 48 hpi (Fig. 1n). Moreover, the DNA load of the *gp41* gene was also significantly reduced at 48 hpi (Fig. 1o). These results implyed that the inhibition of piR-bmo-796514 expression could effectively suppress the proliferation of BmNPV in BmN cells.

The comprehensive results of *in vivo* and *in vitro* experiments indicated that BmNPV infection could induce the expression of piR-bmo-796514 to promote virus proliferation.

### piR-bmo-796514 targets RNF181 to promote the proliferation of BmNPV

In our previous study on the response of piRNAs in the fat body and midgut of silkworm to BmNPV infection, we have analyzed the target genes of the differentially expressed piRNAs ^22^. RNF181 was predicted as a potential target gene of piR-bmo-796514, with a complementary sequence to piR-bmo-796514 in the CDS region (Fig. 2a). RNF181, a member of the E3 ubiquitin ligase family, possesses a typical RING finger conserved domain (Fig. 2b). To determine whether piR-bmo-796514 directly regulates its potential target gene *RNF181*, we first transfected BmN cells with piR-bmo-796514-mimics and quantified RNF181 mRNA levels by qPCR. Notably, RNF181 expression was significantly reduced at 24 and 48 hours post-transfection, implying the inhibition of RNF181 by piR-bmo-796514 (Fig. 2c). Conversely, transfection of BmN cells with piR-bmo-796514-inhibitor resulted in a significant upregulation of RNF181 expression (Fig. 2d). Additionally, we constructed dual-luciferase vectors containing the predicted target sequence of RNF181 or a mutated sequence of the RNF181 target in the CDS region (Supplementary Fig. 3). The dual-luciferase reporter gene assays demonstrated that when BmN cells were co-transfected with the piR-bmo-796514 mimic and the luciferase reporter plasmid containing the wild-type RNF181 target sequence, a significant reduction in luciferase activity was observed (Fig. 2e). However, when the RNF181 target sequence was mutated, this inhibitory effect was abolished (Fig. 2e). In contrast, co-transfection of the piR-bmo-796514 inhibitor and the wild-type RNF181 target sequence led to a significant increase in luciferase activity, and this enhancing effect was eliminated when the RNF181 target sequence was mutated (Fig. 2f). Furthermore, analysis of the expression profile of the target gene *RNF181* by qPCR showed a significant downregulation of RNF181 in both the fat body (24 hpi) and BmN cells (48 hpi) following BmNPV infection (Fig. 2g, i). However, transfection of BmN cells with piR-bmo-796514 inhibitors could rescue the downregulation trend of RNF181 caused by BmNPV infection (Fig. 2h, j), further illustrating the negative regulatory effect of piR-bmo-796514 on RNF181.

**Fig. 2:**
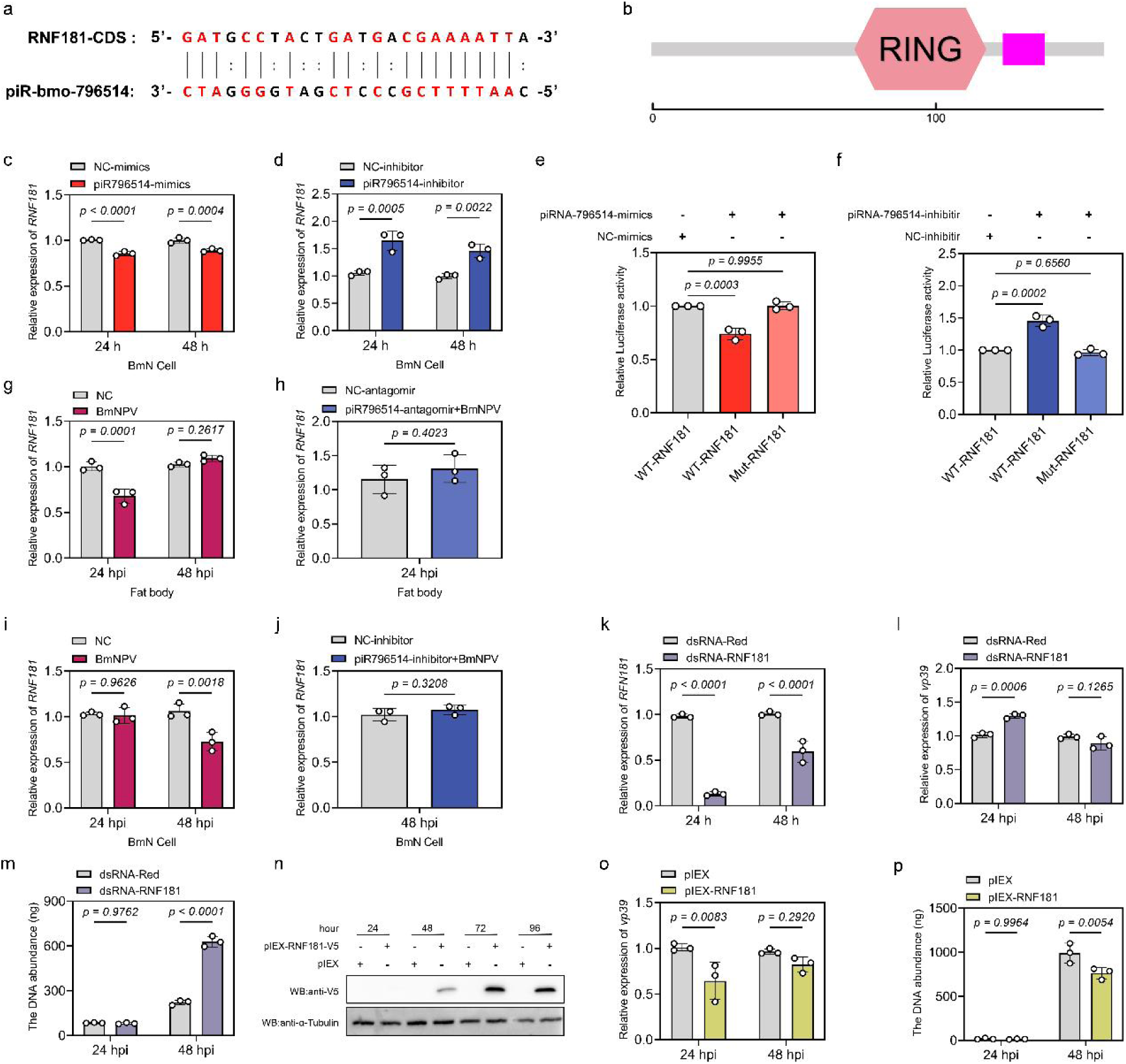
Interaction between piR-bmo-796514 and its target gene RNF181, and the impact of RNF181 on BmNPV proliferation. (a) Sequence complementarity between piR-bmo-796514 and its target gene RNF181. (b) Analysis of the conserved domains of RNF181. (c) Detection of the transcriptional level of the *RNF181* gene in BmN cells after transfection with piR-bmo-796514-mimics. (d) Detection of the transcriptional level of the RNF181 gene in BmN cells after transfection with piR-bmo-796514-inhibitor. (e,f) Dual-luciferase reporter gene assays to test the regulatory effect of piR-bmo-796514 on RNF181 using piRNA-796514 mimics and inhibitor. (g) Quantitative detection of RNF181 transcriptional levels in silkworm fat bodies after BmNPV infection. (h) Quantitative detection of RNF181 transcriptional levels in silkworm fat bodies after injection with piR-bmo-796514-antagomir and subsequent BmNPV infection. (i) Quantitative detection of RNF181 transcriptional levels in BmN cells after BmNPV infection. (j) Quantitative detection of RNF181 transcriptional levels in BmN cells after transfection with piR-bmo-796514-inhibitor and subsequent BmNPV infection. (k) Detection of silencing of RNF181 after transfection with dsRNA-RNF181 in BmN cells. (l) Detection of the transcriptional level of the viral *vp39* gene in BmN cells after transfection with dsRNA-RNF181 and subsequent BmNPV infection. (m) Detection of the DNA load of the viral *gp41* gene in BmN cells after transfection with dsRNA-RNF181 and subsequent BmNPV infection. (n) Western blot detection of RNF181 expression after transfection of BmN cells with pIEX-RNF181-V5. (o) Detection of the transcriptional level of the viral *vp39* gene in BmN cells after overexpression of RNF181 and subsequent BmNPV infection. (p) Detection of the DNA load of the viral *gp41* gene in BmN cells after overexpression of RNF181 and subsequent BmNPV infection. *p*-value <0.05 was considered as statistically significant.

To elucidate how the target gene *RNF181* affected viral infection, we designed specific double-stranded RNA (dsRNA) to knockdown the expression of *RNF181* in BmN cells. The qPCR results of knockdown efficiency revealed that the transcriptional level of *RNF181* was significantly reduced at both 24 and 48 hours post-transfection with dsRNA-RNF181 (Fig. 2k). Moreover, knockdown of *RNF181* in BmN cells and subsequent infection with BmNPV revealed a significant increase in virus *vp39* expression (24 hpi) (Fig. 2l) and viral DNA load (48 hpi) (Fig. 2m).

We further transfected BmN cells with the overexpression vector pIEX-RNF181-V5 or the empty control vector. Western Blot analysis showed that RNF181-V5 was successfully expressed in BmN cells (Fig. 2n). When BmN cells were infected with BmNPV at 24 h after transfection of *RNF181*, it was observed that the mRNA expression level of the viral gene *vp39* (Fig. 2o) and the viral DNA load (Fig. 2p) were significantly down-regulated at 24 or 48 hpi. The results of knockdown and overexpression experiments suggested that RNF181 could inhibit BmNPV proliferation.

### piR-bmo-796514 and RNF181 regulate Integrin α2b-like expression

To explore the exact mechanism by which the piR-bmo-796514 target gene *RNF181* inhibit BmNPV proliferation, the Prediction and Display System for Ubiquitin Ligase-Substrate Interactions (UbiBrowser 2.0)^41^ was employed to predict potential substrate proteins for RNF181. The prediction results revealed multiple candidate substrates for RNF181 in humans, including Integrin αIIb (ITGA2B) (Fig. 3a, Fig. Supplementary 4a, b). Previous studies in human platelets have demonstrated that Integrin αIIbβ3 serves as a substrate for RNF181, with the E3 ubiquitin ligase specifically interacting with the KVGFFKR motif of this integrin^42^. Intriguingly, a homologous integrin family member, Integrin α2b-like, was identified in *Bombyx mori*, sharing a conserved Int α domain with its human counterpart, *Homo sapiens* Integrin α2b (Supplementary Fig. 4c). Further comparative analysis revealed a similarly conserved motif, KIGFFNR, within the amino acid sequence of *B. mori* Integrin α2b-like, suggesting a potential interaction between this integrin and RNF181 in silkworm (Supplementary Fig. 4d). These findings provided a strong foundation for exploring the role of RNF181 in ubiquitin-mediated regulation of Integrin α2b-like and its implications for BmNPV proliferation.

**Fig. 3:**
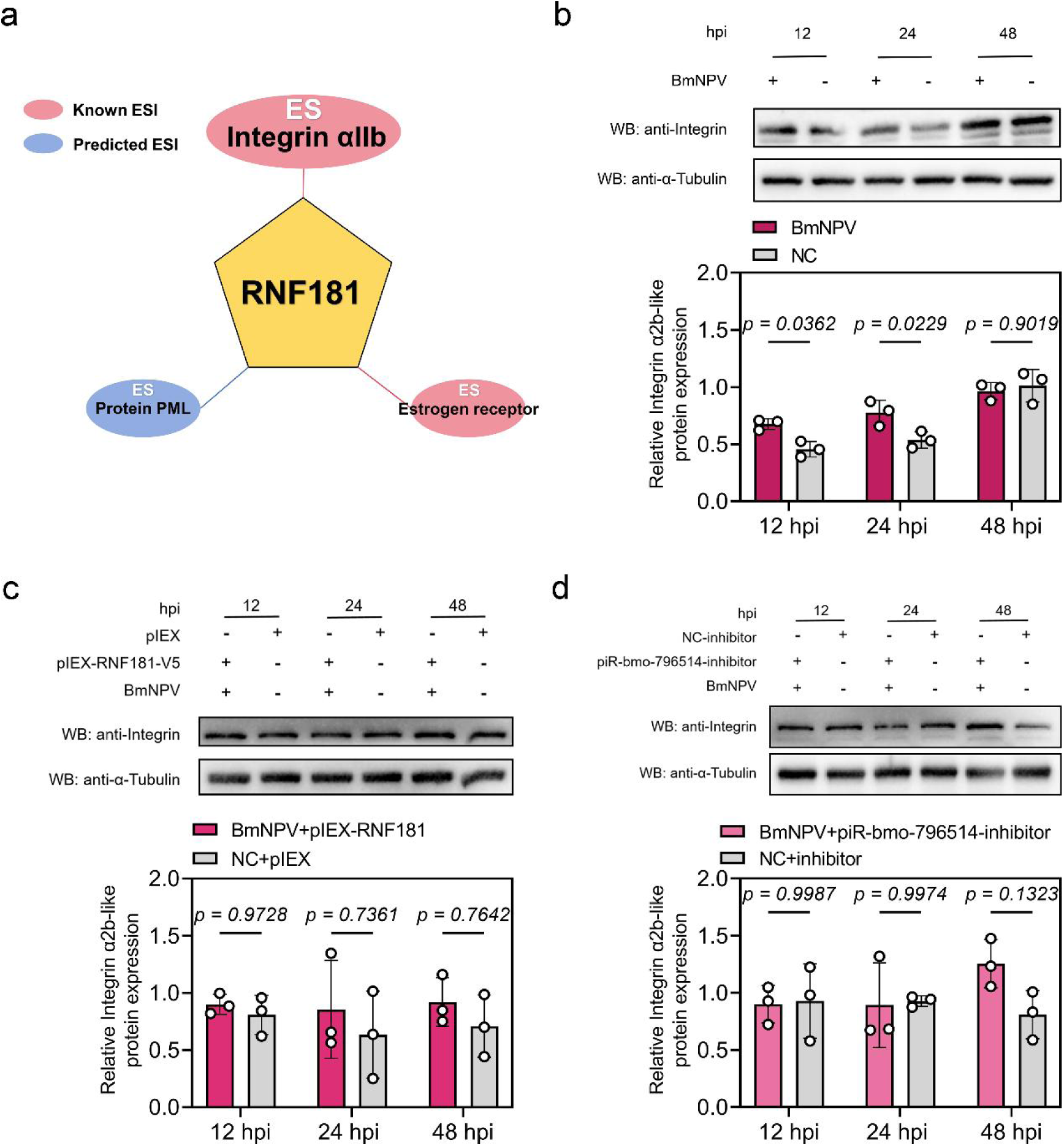
Integrin α2b-like, a potential substrate protein of RNF181, was significantly downregulated after BmNPV infection. (a) Prediction of substrate proteins for RNF181 using UbiBrowser 2.0. (b) Detection of Integrin α2b-like expression levels in BmN cells after BmNPV infection by Western blot. (c) Detection of Integrin α2b-like expression levels in BmN cells after overexpression of RNF181 and BmNPV infection by Western blot. (d) Detection of Integrin α2b-like expression levels in BmN cells transfected with piR-bmo-796514-inhibitor and infected with BmNPV by Western blot. *p*-value <0.05 was considered as statistically significant.

A specific antibody for Integrin α2b-like was used to detect the protein expression levels of endogenous Integrin α2b-like in BmN cells in the presence or absence of BmNPV infection. The results showed a significant increase in Integrin α2b-like protein expression levels at 12 and 24 hpi (Fig. 3b). However, overexpression of RNF181 (Fig. 3c) or treatment with piR-bmo-796514 inhibitor (Fig. 3d) in BmN cells prevented the induction of Integrin α2b-like expression by BmNPV infection at 12 and 24 hpi (Fig. 3c, d).

Collectively, these results suggested that both piR-bmo-796514 and RNF181 were involved in the regulation of Integrin α2b-like expression during BmNPV infection.

### RNF181 interacts with Integrin-α2b-like and facilitates its degradation through ubiquitination

To further determine whether RNF181 interacts with Integrin α2b-like proteins, laser confocal microscopy experiment was employed to investigate the intracellular localization of RNF181 and Integrin α2b-like. Confocal imaging results revealed that the red fluorescence (RNF181) overlapped with the green fluorescence (Integrin α2b-like), indicating co-localization of RNF181 and Integrin α2b-like within the cells (Fig. 4a-c). We further validated the interaction between RNF181 and Integrin α2b-like proteins using Co-Immunoprecipitation (Co-IP). BmN cells were co-transfected with pIEX-Integrin α2b-like-His and pIEX-RNF181-V5, and cell protein samples were collected 72 hours post-transfection for Co-IP experiments. Cell lysates were immunoprecipitated using an anti-His antibody, followed by Western blot analysis to detect pIEX-Integrin α2b-like-His and pIEX-RNF181-V5. The results demonstrated that Integrin α2b-like-His co-precipitated with RNF181-V5, indicating a physical interaction between the two proteins (Fig. 4d). Conversely, reciprocal co-IP experiments using an anti-V5 antibody yielded consistent results, further confirming the association between Integrin α2b-like and RNF181 (Fig. 4e).

**Fig. 4:**
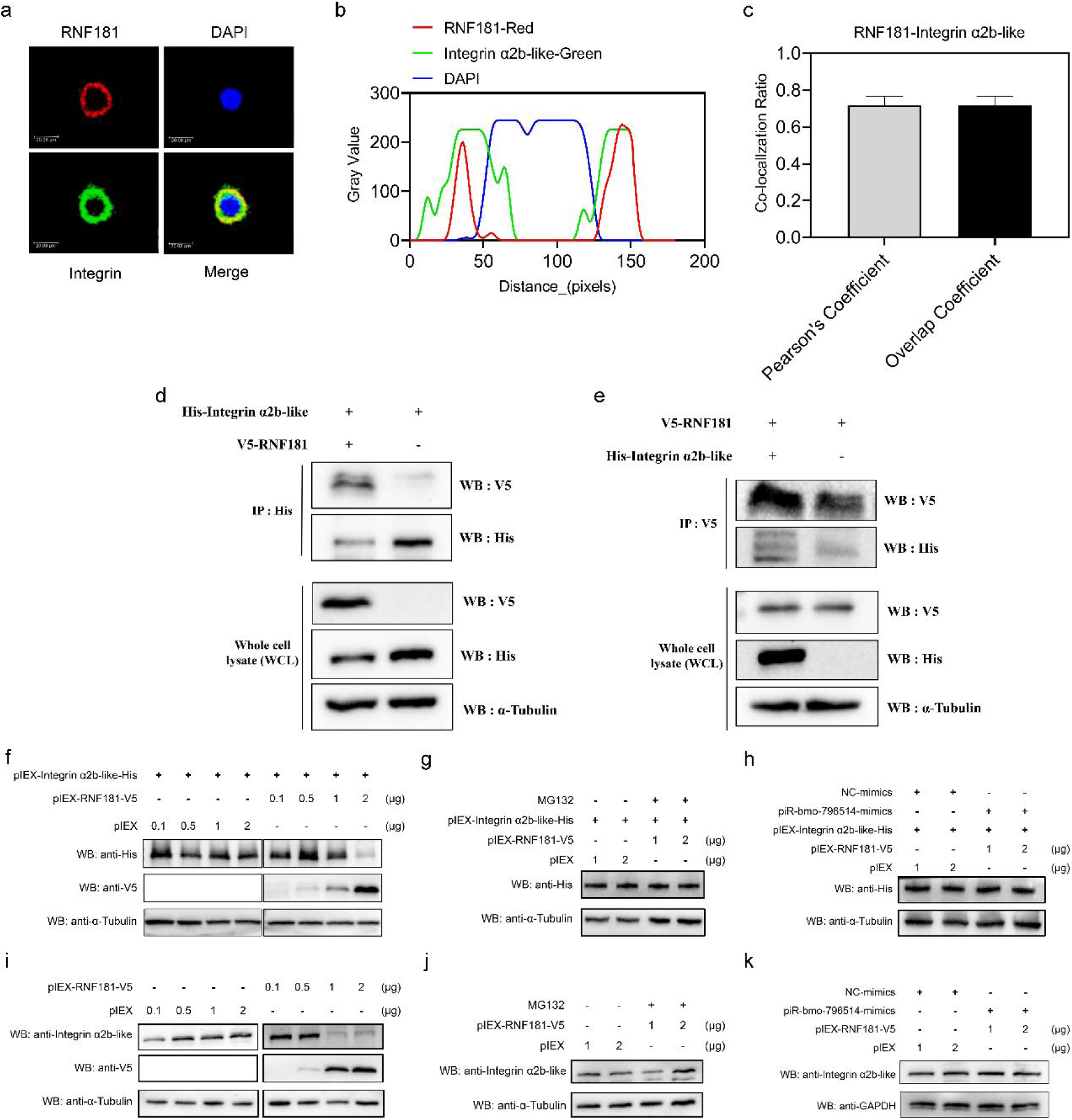
Verification of the interaction between RNF181 and Integrin α2b-like. (a) Immunofluorescence detection of the cellular localization of RNF181 (red) and Integrin α2b-like (green). (b) Analysis of the fluorescence signals of RNF181 and Integrin α2b-like by confocal microscopy scanning. (c) Pearson’s and overlap coefficients of RNF181 and Integrin α2b-like were analyzed using Image J software as described previously ^90,91^. (d, e) Immunoprecipitation assays to detect the interaction between RNF181 and Integrin α2b-like. (f) Western blot detection of the expression levels of exogenous Integrin α2b-like after co-expression with RNF181. (g) Western blot detection of the expression levels of exogenous Integrin α2b-like after co-expression with RNF181 and treatment with MG132. (h) Western blot detection of the expression levels of exogenous Integrin α2b-like after co-expression with piR-bmo-796514 mimics and RNF181. (i) Western blot detection of the expression levels of endogenous Integrin α2b-like after overexpression of RNF181. (j) Western blot detection of the expression levels of endogenous Integrin α2b-like after overexpression of RNF181 and treatment with MG132. (k) Western blot detection of the expression levels of endogenous Integrin α2b-like after co-expression with piR-bmo-796514 mimics and RNF181.

Considering that RNF181 belongs to the family of E3 ubiquitin ligases, the subsequent focus of our research was to investigate whether RNF181 could facilitate the degradation of Integrin α2b-like. We first examined the impact of RNF181 on the expression of exogenous Integrin α2b-like proteins by co-transfecting BmN cells with a fixed concentration of the pIEX-Integrin α2b-like-His and varying concentrations of the pIEX-RNF181-V5 plasmid. The results indicated that RNF181 (2 μg) could promote the degradation of exogenous Integrin α2b-like proteins (Fig. 4f). Furthermore, MG132, a specific inhibitor of the ubiquitin-proteasome pathway, could reverse the degradation of exogenous expressed Integrin α2b-like induced by overexpression of RNF181 (Fig. 4g), indicating that RNF181 degraded exogenous Integrin α2b-like through the ubiquitin-proteasome pathway.

More important, to further elucidate whether piR-bmo-796514 participated in the regulation of Integrin α2b-like degradation, during the co-transfection of RNF181 and Integrin α2b-like, piR-bmo-796514 mimic was also transfected concurrently to downregulate the expression of RNF181. The results indicated that the piR-bmo-796514 mimics could also reverse the degradation of exogenous Integrin α2b-like mediated by overexpression of RNF181 (Fig. 4h).

We further used a specific antibody targeting Integrin α2b-like to detect the degradation of endogenous Integrin α2b-like by RNF181. The results demonstrated that overexpression of RNF181 (1 and 2 μg) could promote the degradation of endogenous Integrin α2b-like (Fig. 4i). Moreover, the treatment with MG132 or piR-bmo-796514-mimics was capable of impeding the degradation of endogenous Integrin α2b-like (Fig. 4j, k). Collectively, these results suggested that RNF181 could degrade Integrin α2b-like via the ubiquitin-proteasome pathway, while piR-bmo-796514 could antagonize the degradation of integrin α 2b like protein by targeting RNF181.

### Integrin-α2b-like establishes an interaction with the GP64 protein of BmNPV to **influence BmNPV invasion.**

To further ascertain the impact of Integrin α2b-like on the proliferation of BmNPV, *Integrin α 2b like* gene was knocked down and overexpressed respectively in BmN cells followed by viral infection experiments.

The results of knockdown efficiency revealed that the transcriptional level of *Integrin α2b-like* was significantly decreased at both 24 and 48 hours post-transfection with dsRNA-Integrin α2b-like (Fig. 5a). Knockdown of *Integrin α2b-like* in BmN cells and subsequent infection with BmNPV revealed a significant decrease in virus *vp39* expression (24 and 48 hpi) and viral DNA load (48 hpi) (Fig. 5b, c). These results suggested that the knockdown of Integrin α2b-like could inhibit the proliferation of BmNPV. Moreover, Western Blot analysis demonstrated the successful expression of the Integrin-his recombinant protein at 24, 48, 72, and 96 hours post-transfection (Fig. 5d). When BmN cells were infected with BmNPV at 24 h after transfection of pIEX-Integrin α2b-like-his, it was observed that the mRNA expression level of the viral gene *vp39* (24 and 48 hpi) (Fig. 5e) and the viral DNA load (48 hpi) (Fig. 5f) were both significantly up-regulated. These findings suggested that Integrin α2b-like could promote the proliferation of BmNPV.

**Fig. 5:**
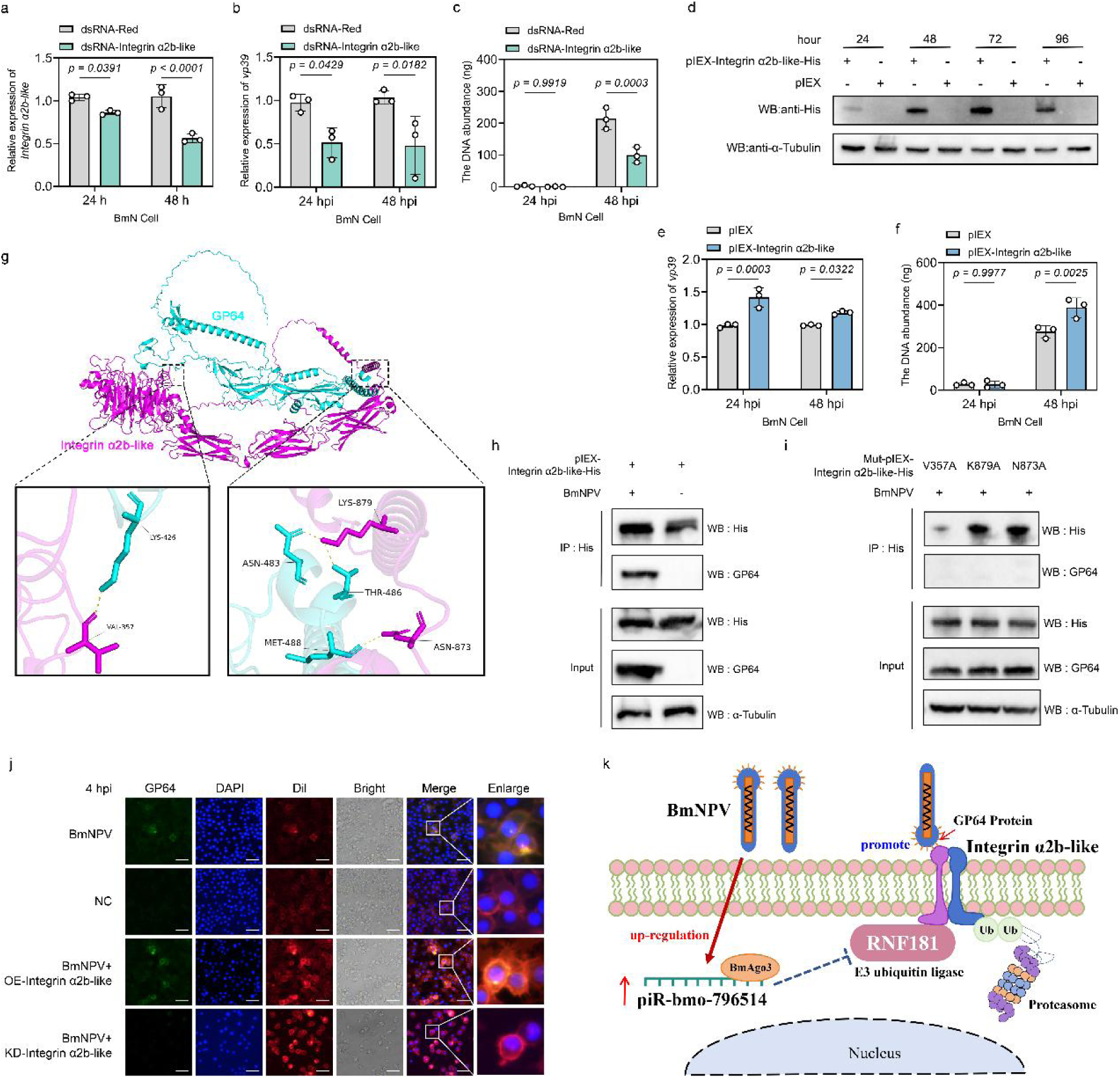
Integrin α2b-like affects the cell entry of BmNPV by binding to the viral envelope protein GP64. (a) Assessment of the silencing of Integrin α2b-like in BmN cells following transfection with dsRNA-Integrin α2b-like. Detection of the transcriptional level of the viral *vp39* gene (b) and DNA load of the viral *gp41* gene (c) in BmN cells that were transfected with dsRNA-Integrin α2b-like and infected with BmNPV. (d) Western blot detection of Integrin α2b-like expression in BmN cells after transfection with pIEX-Integrin α2b-like-His. (e) Detection of the transcriptional level of the viral *vp39* gene and DNA load of the viral *gp41* gene (f) in BmN cells that overexpressed Integrin α2b-like and were infected with BmNPV. (g) Prediction of the binding sites between Integrin αIIb-like and GP64 by AlphaFold3. The structure of the Integrin α2b-like and GP64 protein was obtained from AlphaFold3 predictions. Overall protein structures are shown together with details of the interaction. (h) Immunoprecipitation assay was used to detect the interaction between Integrin α2b-like and the BmNPV envelope protein GP64. (i) Immunoprecipitation assay was employed to detect the interaction between Integrin α2b-like mutants and the BmNPV envelope protein GP64. (j) After BmNPV infection of BmN cells, immunofluorescence was used to detrect viral GP64 expression in BmN cells after overexpression or knock-down of Integrin α2b-like. (k) Schematic of the mechanism of promotion of BmNPV proliferation by piR-bmo-796514. *p*-value <0.05 was considered as statistically significant.

According to the reported binding sites of integrin protein interacting with other proteins ^43^, we identified multiple previously-reported integrin-binding motifs within the amino-acid sequence of BmGP64, such as NVR (Asn-Val-Arg), LDI (Leu-Asp-Ile), and LDA (Leu-Asp-Ala) (Supplementary Fig. 5a). Moreover, the amino-acid sequence of *Bombyx mori* Integrin α2b-like was also found to contain the NVR binding motif, which may be involved in the interaction with other integrins (Supplementary Fig. 5b). An EGR motif in the fusion loop of the postfusion structure of GP64 was also proposed as an integrin-interacting motif^44^.

To further investigate whether GP64 can interact with Integrin α2b-like, we performed structural modeling using AlphaFold3. The protein sequences of GP64 and Integrin α2b-like were submitted to AlphaFold3 for structural prediction, yielding their interaction complex, which was subsequently visualized using PyMOL to identify the interaction sites. The docking analysis revealed multiple putative binding sites including Val-357, Lys-879, Asn-873 at their interface, suggesting a stable molecular interaction (Fig. 5g). To investigate whether Integrin α2b-like interacts with GP64, BmN cells overexpressing Integrin α2b-like were infected with BmNPV, and cell lysates were harvested at 72 hpi. Immunoprecipitation using an anti-His antibody followed by Western blot analysis revealed that Integrin α2b-like-His co-precipitated with GP64, demonstrating an interaction between the two proteins (Fig. 5h). We constructed three Integrin α2b-like mutants, each featuring Aalanine substitutions at key residues targeting the predicted binding sites with GP64. Co-IP assays demonstrated that all three mutants completely lost their ability to bind GP64, confirming the critical role of these residues in mediating the protein-protein interaction (Fig. 5i). The NVR-binding motif in Integrin α2b-like is located in the central region of its three-dimensional structure, which differs from our previous prediction of binding sites at both termini (Fig. S5c). Site-directed mutagenesis was also performed on the NVR-binding motif in integrin α2b-like, substituting NVR with Aalanine residues. Co-IP analysis revealed that the mutant protein failed to interact with GP64, indicating the essential role of this motif in mediating the protein-protein interaction (Supplementary Fig. 5d). Subsequently, we downregulated and overexpressed Integrin α2b-like in BmN cells and then infected these cells with BmNPV at a MOI of 5. Fluorescence microscopy imaging was employed to observe the localization of the viral GP64 protein at 4 hours post-virus inoculation. In the early invasion stage of infection, the result showed that BmN cells incubated with the BmNPV for 4 hours exhibited weak green fluorescence, while cells not incubated with the virus showed no green fluorescence (Fig. 5j). Intriguingly, the green fluorescence from GP64 was distinctly visible within the cells overexpressing Integrin α2b-like (Fig. 5j). In contrast, no GP64 green fluorescence was detected after the knockdown of Integrin α2b-like (Fig. 5j). These findings suggest that during BmNPV infection, Integrin α2b-like interacts with the viral envelope protein GP64, functioning to mediate viral entry.

## Discussion

The first and main function of piRNAs that was discovered was the silencing of transposable elements, which is crucial for maintaining the integrity of the germ line genome^4,6^. Naturally, piRNAs were subsequently found to defend against retroviruses, i.e. RNA viruses that can integrate into the host genome and have a close evolutionary relationship with transposons, in domestic chickens and koalas^45,46^. With respect to exogenous viruses, it was found that the piRNA pathway is not required for antiviral defense in fruit flies ^8^ while it is implicated in antiviral defense in arbovirus (RNA virus)-infected mosquitoes and their derived cell lines^9,12,13,47^. However, in culicine mosquitoes, the antiviral function of the piRNA pathway is considered a consequence of the evolution and expansion of specific PIWI class proteins^48,49^ and it remains controversial to extend this function of the piRNA pathway to other insects. Although virus-derived piRNAs (vpiRNAs) have been identified in mosquito cell lines and in adult *Aedes* mosquitoes infected with various RNA viruses^9,50,51^, no direct evidence exists so far that vpiRNAs have antiviral activity^13,48,49^. While the majority of reports argue for an antiviral role of piRNAs in insects, it is intriguing that our recent study found that the piRNA pathway in silkworms has the potential to promote the proliferation of exogenous DNA viruses^23^. In our research model of BmNPV infection in silkworms, no vpiRNAs were detected, but significant changes in the expression of host derived piRNAs were observed after viral infection^22^. However, there is also no direct evidence to suggest that host derived piRNAs promote the proliferation of exogenous viruses^22,23^. For instance, some studies have preliminarily screened host piRNAs that could target SARS-CoV-2, suggesting that these piRNAs might affect SARS-CoV-2 infection^52–54^. In addition, strong responses in the levels of cellular (host) piRNAs were also identified following Seneca virus A^55^, herpes simplex virus type 1 (HSV-1)^55^, and White Spot Syndrome Virus (WSSV) infection^56^. Most notably, there have been attempts to investigate the impact of specific host derived piRNAs on exogenous viral infections (WSSV)^56^. A recent study discovered that piR-pva-926938, a downregulated piRNA targeting the WSSV186 gene, paradoxically increased viral load by downregulating host immune genes like calcineurin B and dynamin-binding protein^56^. Nevertheless, there have been no reports on the exact mechanisms by which individual vpiRNAs or host-derived piRNAs influence viral infection. Thus, future research should develop new hypotheses and experimental systems to characterize the functions of vpiRNAs and host derived piRNAs in the interaction between exogenous viruses and hosts.

In the silkworm-BmNPV infection system, it was found that piR-bmo-796514, a host-derived piRNA, could be significantly induced in larval fat body and BmN cells and could target and inhibit the expression of E3 ubiquitin ligase RNF181. RNF181 targets the membrane receptor Integrin α2b-like for ubiquitination and proteasomal degradation, which prevents its interaction with the BmNPV envelope fusion protein GP64 that promotes BmNPV entry. In the presence of high levels of piR-bmo-796514, RNF181 expression is suppressed, rescuing integrin α2b-like from degradation and promoting binding of GP64 (Fig. 5k). Intriguingly, piR-bmo-796514 was found to not belong to any piRNA cluster, but may come from five specific potential loci in the silkworm genome, four of which have not yet been annotated, and one corresponding to a gene (BMSK0002231) that is not responsive to BmNPV infection (Supplementary Fig. 1). Unfortunately, it remains unclear how BmNPV infection induced the expression of piR-bmo-796514. During BmNPV infection, expression levels of BmAgo3 and Siwi are modulated^23^ while also the ping-pong cycle seems to be maintained^22^. However, no ping-pong signature was found in piRBase for piR-bmo-796514, which may indicate that its main production occurs by the primary processing pathway^57^. On the other hand, it is known that baculovirus replication in the nucleus is accompanied by chromatin modifications and changes in the chromatin state throughout the cellular host genome^58^, which could result in an increase of transcriptional activity at a particular locus that harbors piR-bmo-796514-like sequences. Following the processing of piR-bmo-796514 precursor transcripts and interaction with target sequences upon BmNPV infection, a positive feedback loop may be established for increased piR-bmo-796514 production at the genetic locus. Such scenario of de novo production of piRNAs has been documented for other instances of environmental stress, such as increased environmental temperature and heat-shock, and is correlated with the presence of inserted TEs^59,60^. However, such hypothesis may be difficult to verify for the animals used in our study because of strain-specific polymorphisms in the structure of piRNA loci and clusters that are commonly observed^59^.

In addition to silencing TEs, the piRNA pathway and piRNAs have also been known to regulate protein-coding genes in various biological processes^4^. For instance, TE piRNAs from *D. melanogaster* embryos were shown to guide Aub to *nanos* mRNA, which encodes a posterior determinant, resulting in the induction of its deadenylation and decay through interaction with the RNA-binding protein Smaug and the CCR4–NOT deadenylation complex^61^. The outcome for mRNA targets depends on the level of base-pairing with guide piRNAs, with higher complementarity being required for slicing^4,62–64^. However, in the absence of slicing, PIWI proteins may act as mere RNA-binding proteins, able to recruit different machineries for mRNA regulation^4^. In our study, hybridization of piR-bmo-796514 to the target sequence of RNF181 is partial and may not induce mRNA slicing. A limitation of our work concerns the elucidation of the mechanism by which piR-bmo-796514 inhibits RNF181 expression. Indeed, further understanding of the molecular mechanisms of piRNA-dependent mRNA regulation will be an exciting challenge for future studies in the field^4^. In future research, we aim to examine how piR-bmo-796514 targets the degradation of RNF181. Key experiments will involve cross-linking immunoprecipitation (CLIP) methods^65^ that will reveal the composition of the complexes associated with piR-bmo-796514 and its targets.

RNF181, a member of the RING protein family, possesses the typical RING domain characteristic of E3 ubiquitin ligases. This family is well-recognized for its role in the regulation of protein dimerization and protein-protein interactions through its ubiquitin ligase activity^66,67^. Current research indicates that RNF181 can regulate multiple signal transduction pathways and impact various types of cancer in humans. For instance, RNF181 can modulate signaling pathways that involve Hippo/YAP, CARD11 and ERK/MAPK-cyclin D1/CDK4, and it has been implicated in diseases like triple negative breast cancer and gastric cancer^67–71^. Our findings reveal that RNF181 functions to inhibit BmNPV replication in silkworm BmN cells (Fig. 2). Furthermore, we hypothesize that RNF181 exerts its function potentially through its E3 ubiquitin ligase activity, which targets substrate proteins for ubiquitination and degradation. Through prediction in silico, we identified Integrin α2b-like as a potential substrate protein of RNF181 (Fig. 2a. Supplementary Fig. 4). Examination of Integrin α2b-like protein expression levels post-BmNPV infection showed a significant upregulation (Fig. 2b). To verify whether the upregulation of Integrin α2b-like is regulated by RNF181 and piR-bmo-796514, we over-expressed RNF181 or transfected with piR-bmo-796514 inhibitors in the context of BmNPV infection and found a suppression of the upregulation of Integrin α2b-like (Fig. 2c, d). It has also been reported that RNF181, as a novel E3 ubiquitin ligase, interacts with the KVGFFKR motif of platelet integrin αIIbβ3^42^. Combining our interaction experiments between RNF181 and Integrin α2b-like, along with the observation of degradation of Integrin α2b-like by RNF181 (Fig. 4), our results suggest that the expression of Integrin α2b-like is regulated by the upstream factors RNF181 and piR-bmo-796514.

Our immunofluorescence results demonstrate that expression levels of Integrin α2b-like correlate with increased viral gene expression and replication (Fig. 5j), highlighting the significant role of Integrin α2b-like in the process of BmNPV cell entry. Our Co-IP experimental results also confirmed the interaction between Integrin α2b-like and the BmNPV GP64 envelope protein (Fig. 5g-i). Thus, through the analysis of downstream targets of the differentially expressed piR-bmo-796514, our study potentially provided new insights into the cell entry mechanism of BmNPV.

Baculovirus entry of host cells can be mediated by both clathrin-dependent endocytosis and macropinocytosis and is triggered by low pH at the cellular membrane^72,73^. Heparin sulfate, phospholipids and the membrane protein *Bombyx mori* receptor expression-enhancing protein (BmREEPa) all have been proposed as host cell receptors for budded virus attachment and binding^74–76^. More recently, cholesterol and the Niemann-Pick C1 and C2 transport proteins have been implicated in the BmNPV entry mechanism^77^. It is well documented that, in class I alphabaculoviruses, GP64, a class III fusion protein of the viral envelope, is both necessary and sufficient for the binding to cellular membranes and triggering membrane fusion at low pH condition^78^. While the identity of the cellular receptor for GP64 remains elusive, an increasing number of viruses has been found to interact with integrins^79^. Integrins are not only key receptors for the binding of numerous viruses to the cell surface, but some studies have also found that integrins are indispensable for subsequent signaling events that support the infection and replication processes of viruses^80–82^. Studies in murine fibroblast cells have shown that αVβ3 Integrin is essential for the replication of various Flaviviruses^83^. Additionally, research has discovered a role for the αV Integrin Subunit in Varicella-Zoster Virus-mediated fusion and infection^84^. Surface proteins of some viruses, such as the capsid protein VP1 of coxsackievirus A9, contain commonly known integrin-binding sequences, such as RGD, that bind integrins to gain entry into cells or modulate downstream signaling pathways^43^. Additionally, research on Herpes simplex virus has discovered that the virus’s Type 2 Glycoprotein H interacts with αVβ3 Integrin, facilitating viral entry^85^. Both integrin αVβ8 and αVβ6 act as receptors for Foot-and-Mouth Disease Virus^86,87^. Furthermore, studies in silkworms have indicated that Integrin beta serves as a receptor involved in the cell entry of Bombyx mori cypovirus^88^. Interaction of GP64 with integrin has been proposed in a previous structural study^44^. Given the increasing evidence of integrins acting as essential membrane proteins to mediate the entry of a variety of viruses, it may not be surprising that also baculoviruses utilize this pathway to increase their infection efficiency.

In conclusion, our study unraveled for the first time the molecular mechanism by which a host-derived piRNA can mediate the immune escape response during infection of an exogenous DNA virus (Fig. 5k). We believe that our work will stimulate additional research on the impact of specific host piRNAs during exogenous virus infection.

## Methods

### Cell culture, silkworm strain and virus

The *B. mori* ovarian cell line (BmN) was cultured at 28℃ in Grace’s medium (Gibco, USA) containing 10% fetal bovine serum (Gibco, USA), 100 U/mL penicillin, and 100 μg/mL streptomycin (Gibco, USA). Silkworm larvae (*B. mori*, Dazao strain) were reared with fresh mulberry leaves (*Morus* sp.) under constant environmental conditions of 28℃ and 60%–70% humidity. Recombinant BmNPV-EGFP (enhanced green fluorescent protein), as reporter viruses, were constructed by the Bac-to-Bac Baculovirus Expression System. The above cell lines, silkworm strain and virus were kept in Guangdong Provincial Key Laboratory of Agro-animal Genomics and Molecular Breeding.

### piRNA mimics, inhibitors, agomirs and antagomirs

piRNA mimics are small RNA molecules designed based on the sequences of mature piRNAs, which are used to simulate the sequences of endogenous mature piRNAs. Mimics consist of a sequence identical to that of the mature piRNA, as well as a sequence complementary to the mature piRNA target sequence. piR-bmo-796514 mimic (piR-bmo-796514-mimic, GenePharma Tech, China) was used to upregulate piR-bmo-796514 expression. piRNA inhibitors are designed as 21-30 nt RNA oligonucleotides with 2’-methoxy modification, which can effectively inhibit the function of endogenous mature piRNAs. A piR-bmo-796514 inhibitor from GenePharma Tech (China) was employed to downregulate the expression of piR-bmo-796514. Meanwhile, a nonspecific scrambled RNA sequence was used as a negative control (inhibitor-NC or mimics-NC). Transfections were performed using FuGENE HD (Promega, USA) according to the manufacturer’s protocol.

Agomirs and antagomirs have identical sequences as mimics and inhibitors, respectively, but carry additional modifications, such as: cholesterol at 3’ end, two and four phosphorothioates at the backbone at the 5’- and 3’-end, respectively, and methoxy groups throughout the sequence. Compared with ordinary piRNA mimics and inhibitors, agomirs and antagomirs are particularly suitable for *in vivo* animal experiments because of higher stability and cell penetration.

In the in vivo experiment on silkworm larvae, piR-bmo-796514-agomir and piR-bmo-796514-antagomir (GenePharma Tech, China) were used to regulate the expression of piR-bmo-796514. piR-bmo-796514-agomir or piR-bmo-796514-antagomir were incubated with the transfection reagent for 15 minutes. A microsyringe was used to inject the mixture into the silkworm larvae through the hind-foot. The sequences of the aforementioned RNAs are provided in Supplementary Table S1.

### Plasmids and transfection

The pIEX vector was employed for overexpression experiments. The RNF181 ORF (GeneID:101743467) with V5-tag sequences were inserted into the pIEX vector via *Nco*I and *Sac*I to construct the overexpression vector pIEX-RNF181-V5. Similarly, the overexpression vector pIEX-Integrin α2b-like-His for Integrin α2b-like (GeneID:101743745) was constructed. The pGL3-Enhancer (Firefly luciferase) and pRL-SV40-N (Renilla luciferase) vectors (Promega) were used for the dual-luciferase reporter assay. The mRNA sequence of the target gene RNF181, including the 5’UTR and 3’UTR, was inserted into the pGL3-Enhancer vector using *Nco*I and *Sac*I. Additionally, the seed recognition region of RNF181 was mutated using a PCR-based mutagenesis method to obtain the pGL3-RNF181-Mut mutant vector. To systematically investigate the molecular interactions, we generated a series of integrin α2b-like mutants using the QuickMutation™ Site-Directed Mutagenesis Kit (Beyotime, China), named pIEX-Integrin α2b-like-V357A-His, pIEX-Integrin α2b-like-N873A-His, pIEX-Integrin α2b-like-K879A-His and pIEX-Integrin α2b-like-N632A/V633A/R634A-His. These mutants contained Alanine substitutions at key residues within the predicted GP64 binding sites. All primer sequences are listed in Table S2.

For transfection, BmN cells were evenly seeded into a 12-well plate, counted to ensure approximately 10^^5^ cells per well, and incubated overnight at 28℃ until the density exceeded 80%. For transfection with plasmids or dsRNA, a pre-mix system of 100 μL per well was prepared under a laminar flow hood. In a sterile PCR tube, 100 μL of Grace serum-free medium, 1.5 μg of plasmid or 5 μg of dsRNA, and 2 μL of transfection reagent were mixed, incubated at room temperature for 15 minutes. After incubation, the transfection pre-mix was added to the cell plate, gently shaken to mix, and then continued to be cultured at 28℃. After 6 hours, the transfection medium was replaced with complete medium.

### RNA immunoprecipitation (RIP) assay

The BmN cells were collected and suspended in lysis buffer (Beyotime, China). The entire lysate was incubated with antibodies against V5 (Thermofisher, USA) or IgG (Beyotime, China). RNA-protein complexes were recovered with protein A/G plus-agarose (Beyotime, China) and then washed with lysis buffer four times. The precipited piRNA was analyzed by qRT-PCR.

### Virus infection

BmN cells were cultured in 12-well plates and allowed to grow to a density of 70-80%. The cells were then infected with the reporter virus BmNPV-EGFP at a multiplicity of infection (MOI) of 1. After incubation at 28℃ for 1 hour, the medium was replaced with fresh growth medium. Cells were harvested for RNA or DNA extraction at 24-72 hours post-infection (hpi). For virus infection of silkworm larvae, larvae on the first day of the fifth instar were selected. A microsyringe was used to inject the reporter virus into the larvae through the hind-foot, and each larva was injected with 5 μL of the virus (1.0 × 10^^7^ 50% tissue infection dose [TCID_50_]/mL). Viral infection was continuously monitored by fluorescence observation. Silkworm larvae injected with H_2_O served as controls. Larvae were dissected at 24 and 48 hpi, and the fat body tissue was collected. Each group consisted of at least three silkworm larvae, with a minimum of three independent experiments conducted.

### RNA extraction, DNA extraction and real-time qRT-PCR

RNA was extracted from BmN cells and fifth-instar silkworm larvae samples to study the expression of cellular and viral mRNA following viral infection. Total RNA was isolated using the NucleoZOL RNA isolation kit (Macherey-Nagel, Germany) according to the manufacturer’s instructions. Complementary DNA (cDNA) was synthesized from the extracted total RNA using the gDNA Eraser reverse transcription kit (Takara, Japan), following the manufacturer’s protocol.

DNA was extracted from BmN cells and fifth-instar silkworm larvae samples to investigate viral DNA replication. Total DNA was extracted using the Universal DNA extraction kit (Agbio, China) as per the manufacturer’s guidelines. All samples were stored at −20℃ until further use.

Real-time polymerase chain reaction (qPCR) was performed on a Bio-Rad CFX96 Real-Time PCR Detection System using the iTaq Universal SYBR Green Supermix (Bio-Rad, USA). The silkworm *tif4a* gene served as an endogenous control, and the expression of the BmNPV capsid protein *vp39* gene was used to assess viral gene expression. Data analysis was conducted using the 2^^-ΔΔCT^ method. Quantification of viral DNA load was based on detection of the BmNPV *gp41* gene. Standard curves were plotted based on previous studies (Vanarsdall et al., 2005). All primer sequences are listed in Table S2.

### Stem-loop qRT-PCR for piRNA-quantification

We employed the standard qPCR detection method for piRNAs as previously described^22^. Total RNA was extracted using the NucleoZOL RNA isolation kit (Macherey-Nagel, Germany) following the manufacturer’s instructions. cDNA specific to piR-bmo-796514 was generated using a stem-loop RT primer in the reverse transcription reaction, with the RT primer sequences provided in Table S1. Quantitative PCR (qPCR) was conducted on the Bio-Rad CFX96 Real-Time PCR Detection System with iTaq™ Universal SYBR^®^ Green SuperMix (Bio-Rad, USA) using specific qPCR primers for piR-bmo-796514. U6 was utilized as an endogenous control, and data analysis was performed using the 2^-ΔΔCT^ method, with primer sequences listed in Table S2.

### Luciferase reporter assay

The firefly luciferase reporter vector pGL3-Enhancer, the renilla luciferase vector pRL-SV40-N (as an internal control), and piRNA mimics or inhibitors were co-transfected into BmN cells, with the mimic-NC or inhibitor-NC serving as a negative control. After incubation at 28℃ for 72 hours, the activity levels of Firefly Luciferase (FLuc) and Renilla Luciferase (RLuc) in BmN cells were measured using the Dual-Luciferase Reporter Assay Kit (Promega, USA), following the manufacturer’s protocol.

### RNA interference and Over-expression

The T7 RiboMAX™ Express RNAi System (Promega, USA) was used to synthesize dsRNA according to the manufacturer’s instructions. The primers for the target genes that contain a T7 RNA polymerase promoter are listed in Supporting Information Table S1. DsRNA (1 μg/well) was transfected into BmN cells and cell samples were collected at 24 and 48 h post-transfection to detect the expression of target genes. Transfection of dsRNA-Red was used as the control.

BmN cells were transfected with either pIEX-RNF181-V5 or pIEX-Integrin α2b-like-His or pIEX (control) at 1 μg/well. Protein samples were collected and prepared at 24, 48, 72, and 96 h after transfection. The expression of RNF181-V5 and Integrin α2b-like-His proteins was detected by Western blotting using mouse anti-V5 antibody (Thermofisher, USA) and mouse anti-His (Proteintech, China). α-Tubulin was also detected as an internal reference.

### Western blot analysis

Cell protein samples were prepared by RIPA lysis buffer (Beyotime, China), separated by sodium dodecyl sulfate-polyacrylamide gel electrophoresis and transferred to polyvinylidenefluoride membranes for incubation with anti-V5 (ThermoFisher, USA), anti-His (Proteintech, China), anti-GP64 (Abcam, UK), anti-Integrin α2b-like (polyclonal antibody produced in mice) or ɑ-tubulin antibodies (Beyotime, China). Horseradish peroxidase-conjugated goat anti-mouse or anti-rabbit (Beyotime, China) was used as secondary antibody. Western blot signals were detected with the ECL Western Blotting Detection System (Bio-Rad, USA).

### Co-immunoprecipitation (Co-IP)

In BmN cells, RNF181-V5 and Integrin α2b-like-His were co-overexpressed. 72 hours post-transfection, cells were washed twice with PBS and lysed with NP40 lysis buffer (Beyotime, China). After incubation on ice for 30 minutes, the lysates were centrifuged at 14,000 g for 5 minutes, and the supernatants were collected. 20 μL of magnetic beads conjugated with His-tag mouse monoclonal antibody (Beyotime, China) were added to the lysates and incubated overnight at 4℃ on a rotating mixer. The supernatant was then removed using a magnetic stand, and the beads were washed three times with TBS buffer. After washing, 100 μL of SDS-PAGE loading buffer was added, and the mixture was heated at 100℃ for 5 minutes in a water bath. Eluted proteins were detected using anti-V5 and anti-His antibodies. Conversely, the Integrin α2b-like was isolated using magnetic beads conjugated with V5-tag mouse monoclonal antibody (Beyotime, China). The same method was employed to investigate the interaction between Integrin α2b-like and GP64.

### Protein structure prediction and docking

The structural interaction between host Integrin α2b-like (NP_001296513) and BmNPV GP64 protein (QWC64763) was predicted using AlphaFold3^89^. The structure of the Integrin α2b-like and GP64 protein was obtained from AlphaFold3 predictions. Prediction confidence metrics were reported as pTM (predicted Template Modeling score) + ipTM (interface pTM) > 0.5, pLDDT (predicted Local Distance Difference Test) > 50 and PAE (Predicted Aligned Error) < 10. Structural visualization and interaction profiling were performed using PyMOL v2.5.4.

### Immunofluorescence microscopy analysis

Co-localization of RNF181 and Integrin α2b-like: BmN cells were seeded into confocal dishes at an appropriate density, and co-transfected with the overexpression plasmids pIEX-RNF181-V5 and pIEX-Integrin α2b-like-His. Rabbit polyclonal anti-V5 and mouse monoclonal anti-His antibodies were used as primary antibodies, followed by incubation with CoraLite594 goat anti-rabbit (Proteintech, China) or CoraLite488 goat anti-mouse (Proteintech, China) as fluorescent secondary antibodies. Cell nuclei were counterstained with DAPI, and images were captured using a Leica confocal microscope.

Detection of Integrin α2b-like’s effect on BmNPV invasion: BmN cells were seeded into confocal dishes and subjected to overexpression and knockdown treatments, followed by infection with BmNPV-EGFP at an MOI of 5. The cells were fixed 4 hours post-infection and incubated with mouse monoclonal anti-GP64 antibody (Abcam, Britain) and CoraLite488 goat anti-mouse fluorescent secondary antibody (Proteintech, China). Cell nuclei were stained with DAPI and membranes were visualized with the lipohilic stain Dil (Beyotime, China). Images were acquired using a Nikon fluorescence microscope.

### Statistical analysis

The experimental data are expressed as the mean ± standard deviation (SD). Data were analyzed using Student’s *t*-test for comparison of 2 groups or two-way analysis of variance for comparison of multiple groups (GraphPad Prism 8.0. GraphPad, San Diego, CA, USA). *p*-value <0.05 was considered as statistically significant. All data were averaged from 3 independent experiments with cells and 3 independent experiments with larvae that are presented as the mean ± SD.

## Statement of Ethics

An ethics statement was not required for this study type, no human or animal subjects or materials were used.

## Author Contributions

JX and SF did the experiments, collected and analyzed data and drafted the manuscript. WL, MZ, YK, and YH helped with sample preparation, some experiments, and data analysis. LS revised the manuscript and participated in the data analysis. MF designed the reseach scheme, collected and analyzed data, drafted and revised the manuscript. All authors read and approved the final manuscript.

## Funding

This work was supported by the National Natural Science Foundation of China (32372945), Guangzhou fundamental research project (2025A04J5442), Specific university discipline construction project (2023B10564003), National Foreign Expert Human Project (H20240495) and South China Agricultural University high-level talent launch project.

## Competing financial interests

The authors declare that they have no conflicts of financial interest.

---

**Supplementary Fig. 1:**
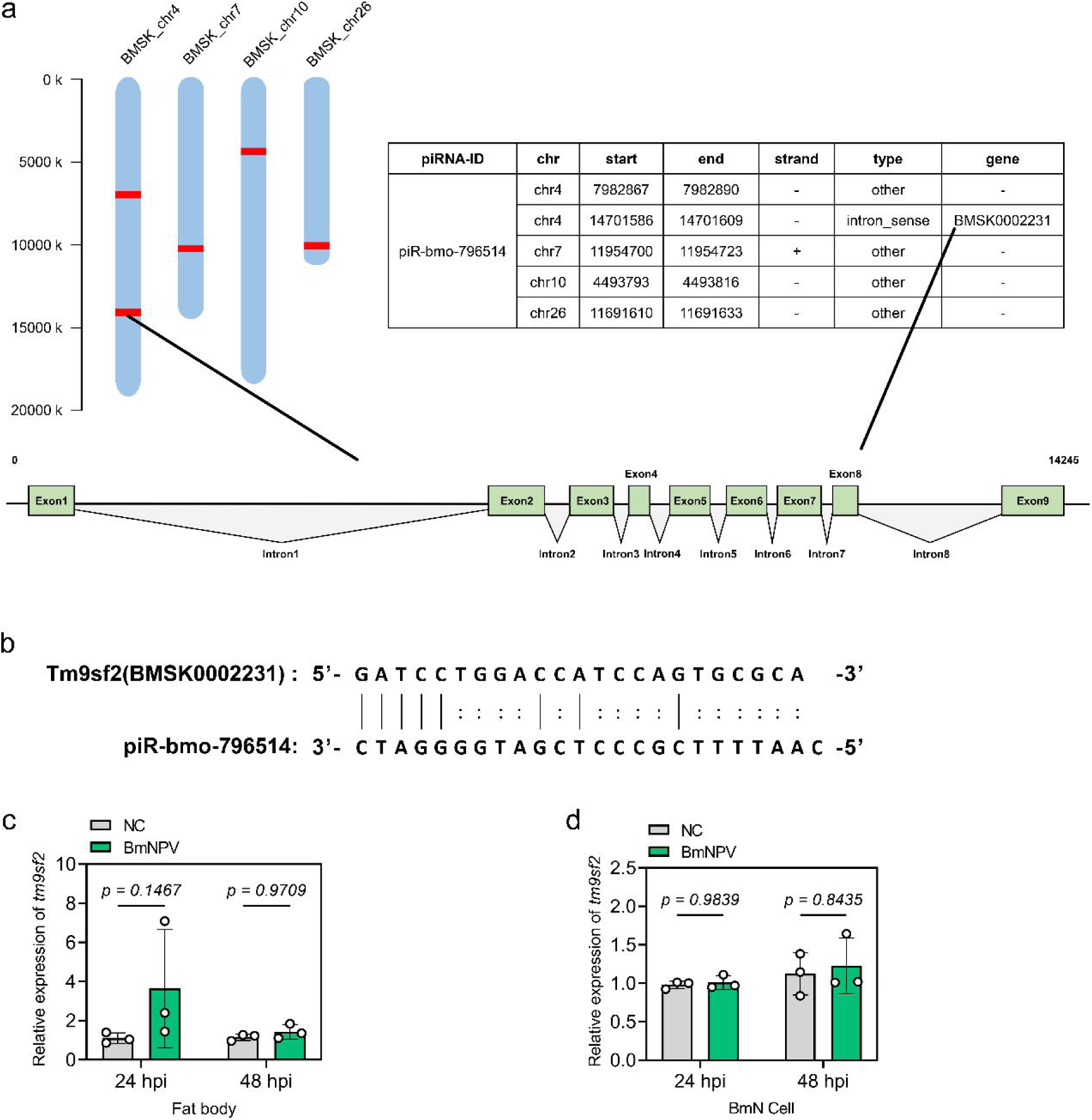
Potential sites of origin of piR-bmo-796514. (a) Potential genomic loci containing piR-bmo-796514 sequence. (b) Sequence complementarity between Tm9sf2(BMSK0002231) and piR-bmo-796514. (c) Quantitative detection of Tm9sf2(BMSK0002231) transcriptional levels in silkworm fat bodies after BmNPV infection. (d) Quantitative detection of Tm9sf2(BMSK0002231) transcriptional levels in BmN cells after BmNPV infection. *p*-value <0.05 was considered as statistically significant.

**Supplementary Fig. 2:**
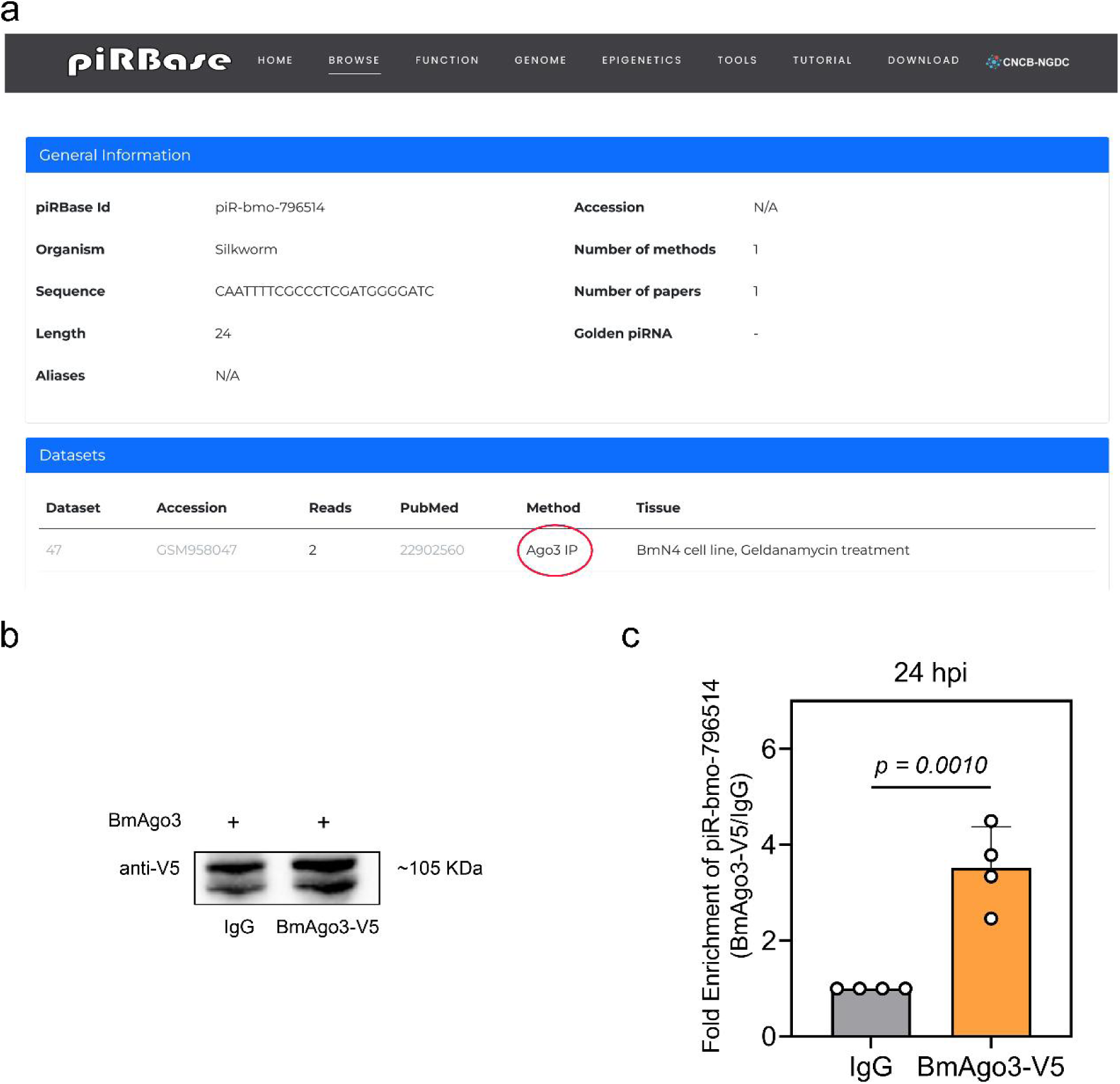
Confirmation of binding of piR-bmo-796514 to BmAgo3. (a) piR-bmo-796514 information in piRBase database. (b) Western blot detection of BmAgo3-V5 expression in BmN cells. (c) RIP analysis of piR-bmo-796514 and BmAgo3 interaction. *p*-value <0.05 was considered as statistically significant.

**Supplementary Fig. 3.**
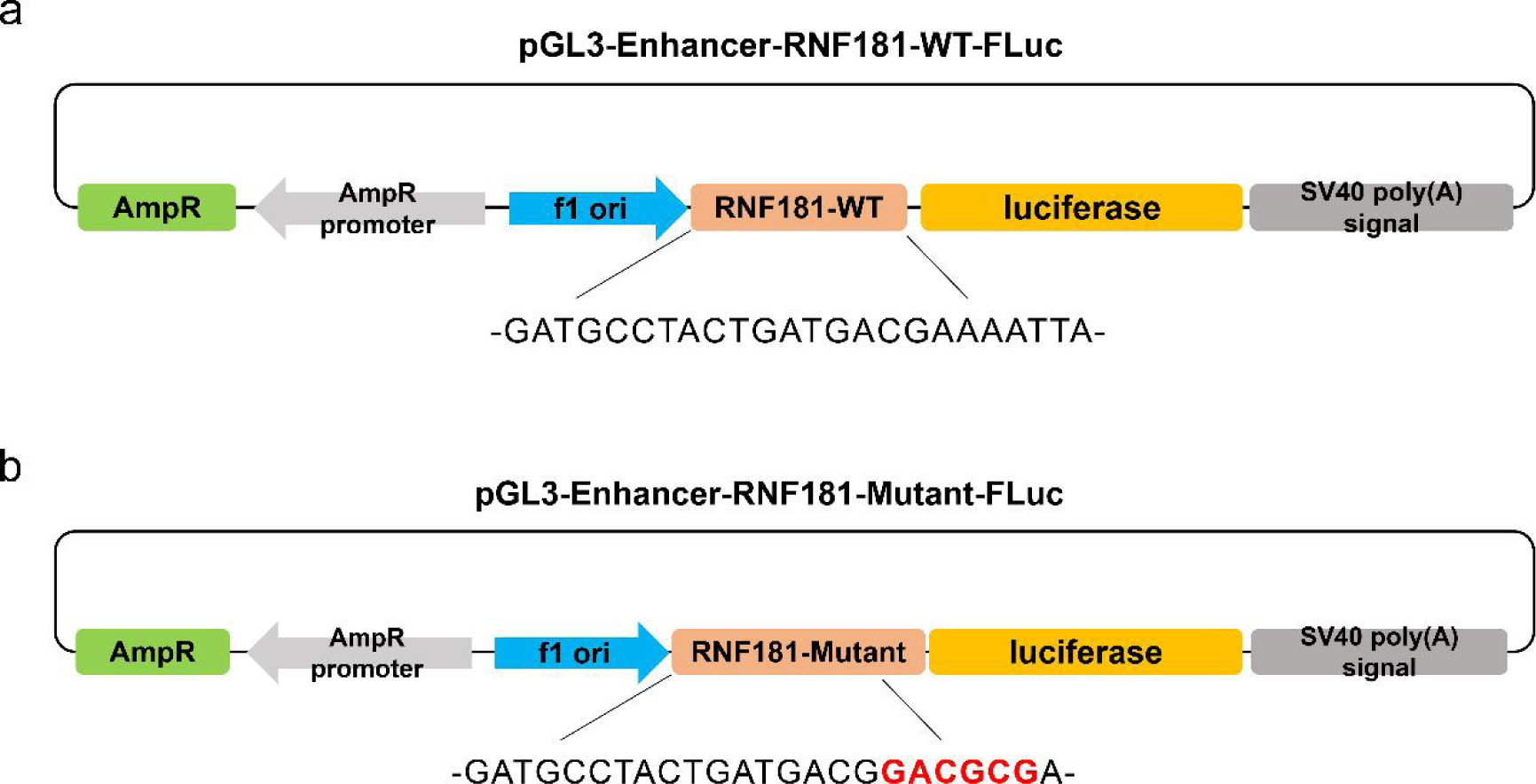
Construction of dual luciferase vectors for RNF181. (a) wild-type RNF181. (b) mutant.

**Supplementary Fig. 4:**
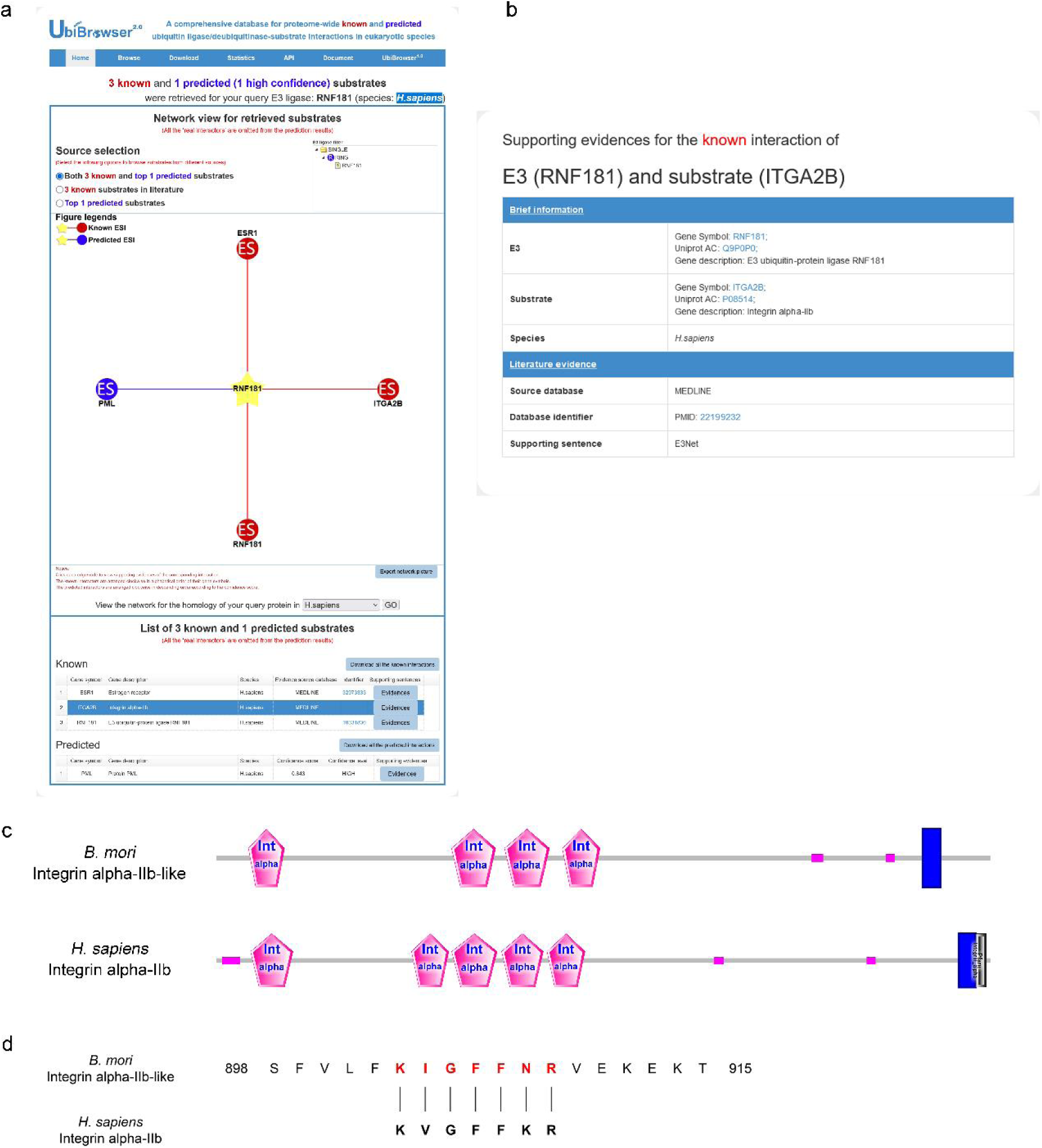
Prediction of RNF181 substrate proteins. (a, b) Utilization of the UbiBrowser 2.0 online tool for the prediction of substrate proteins for human RNF181. (c) Domain analysis of *H. sapiens* Integrin α2b-like in comparison with *B. mori* Integrin α2b-like. (d) Comparison of RNF181 binding motifs within the sequences of *H. sapiens* and *B. mori* Integrin α2b-like.

**Supplementary Fig. 5:**
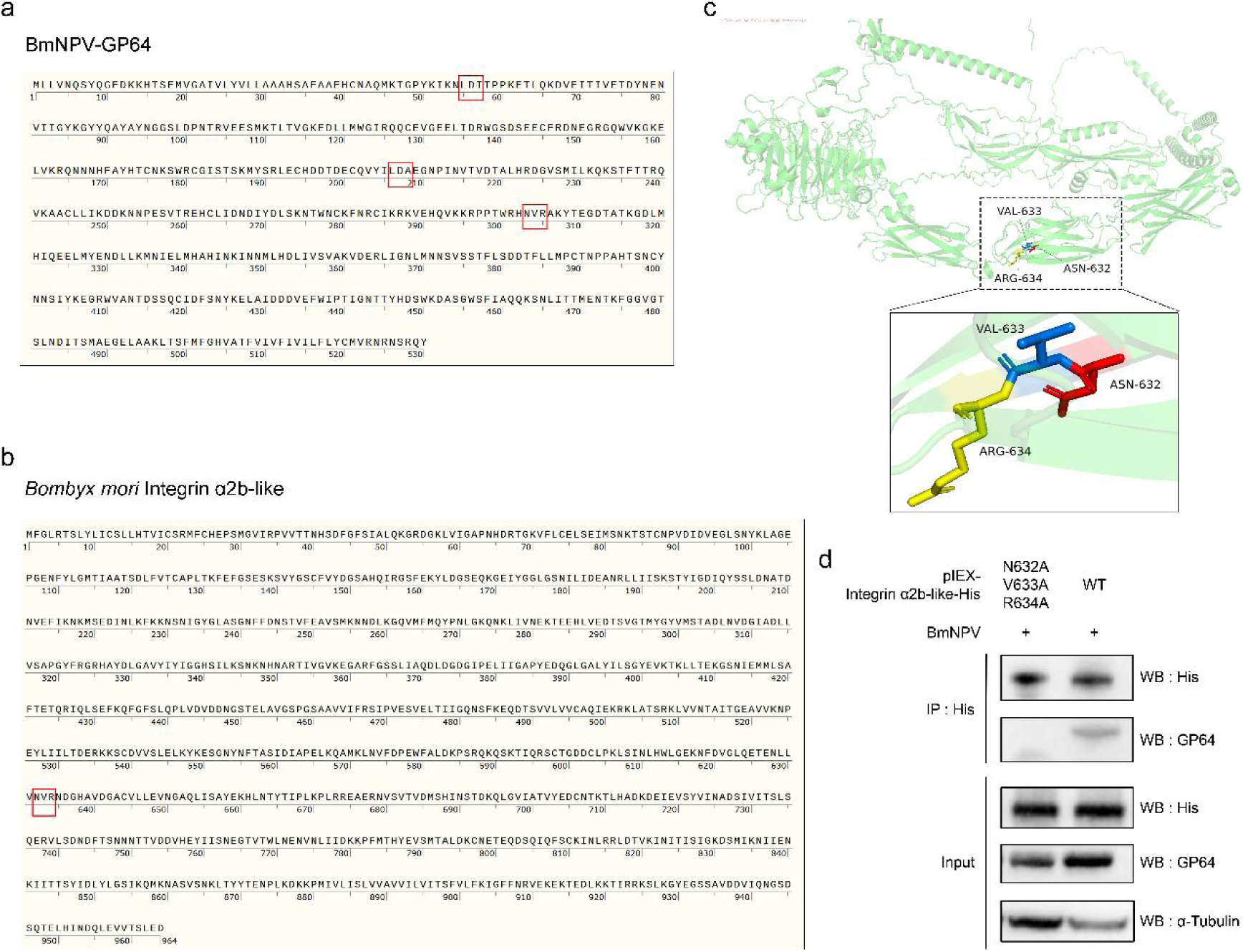
Potential binding motifs in BmNPV envelope protein GP64 and *B. mori* Integrin α2b-like. (a) Potential integrin binding motifs within the GP64 protein. (b) Potential integrin binding motifs within *B. mori* Integrin α2b-like. (c) Immunoprecipitation assay to detect the interaction between Integrin α2b-like mutants and the BmNPV envelope protein GP64.

**Supplementary Table 1:**
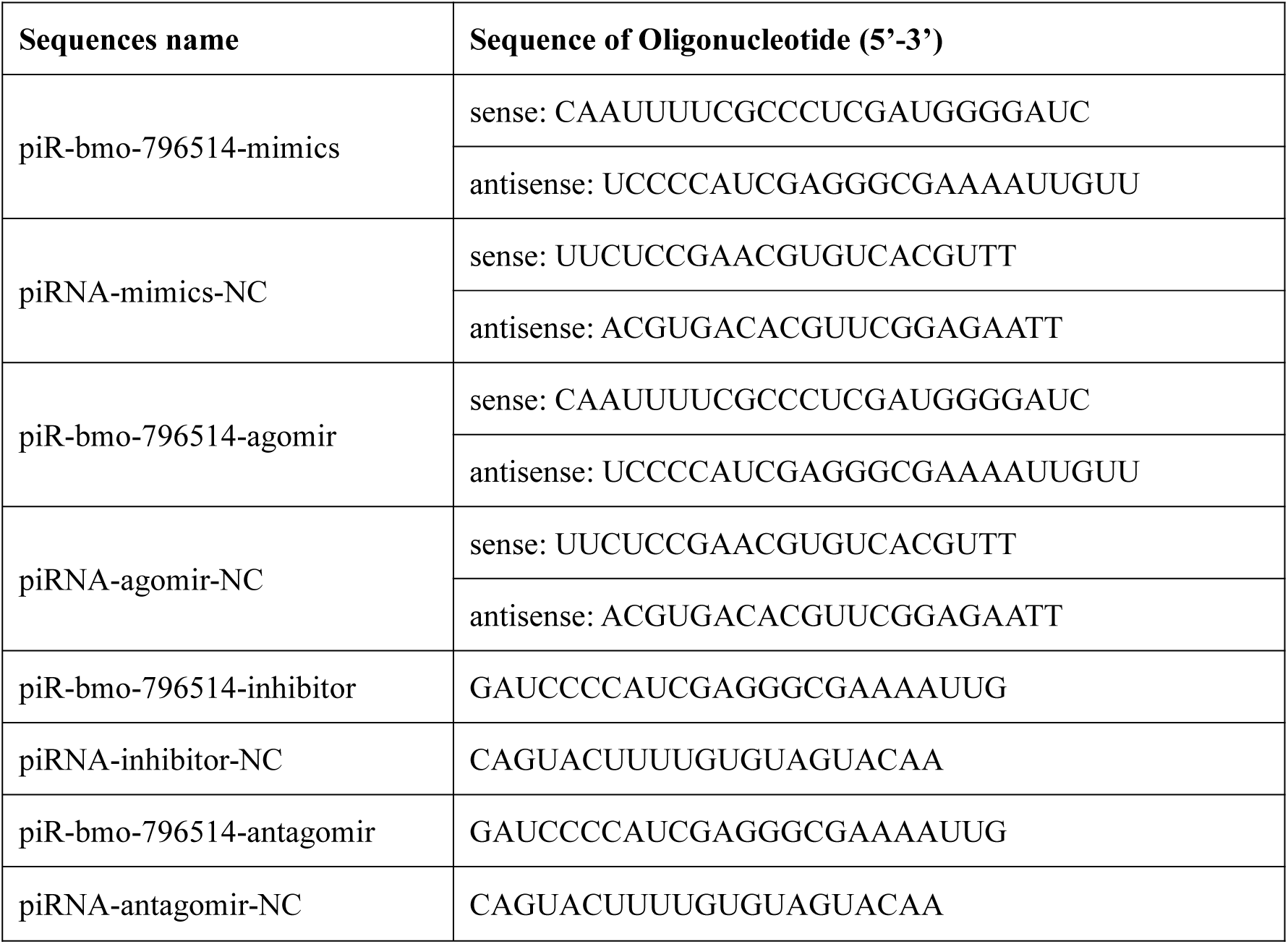
Sequences of piRNA mimics and inhibitors.

**Supplementary Table 2:**
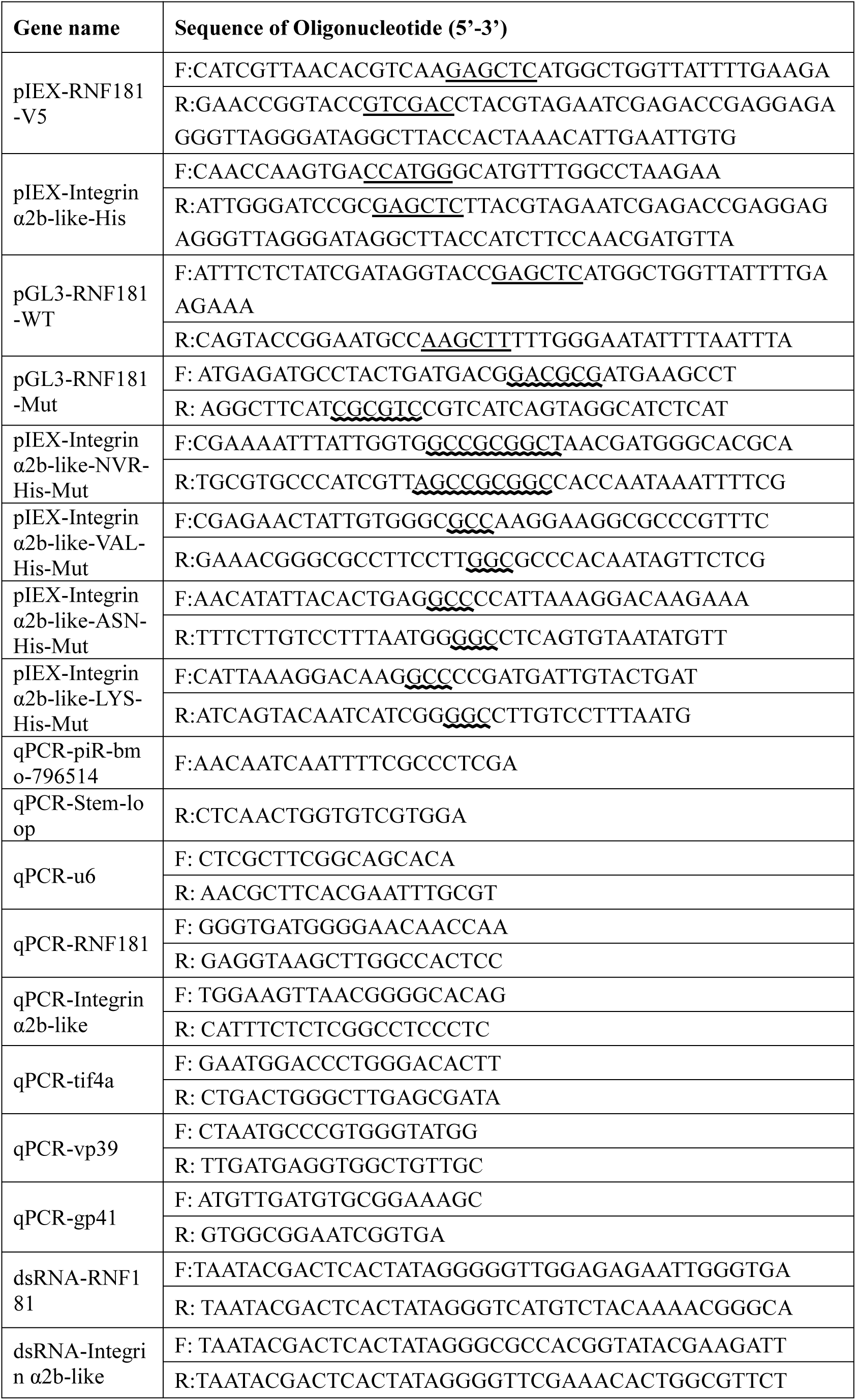
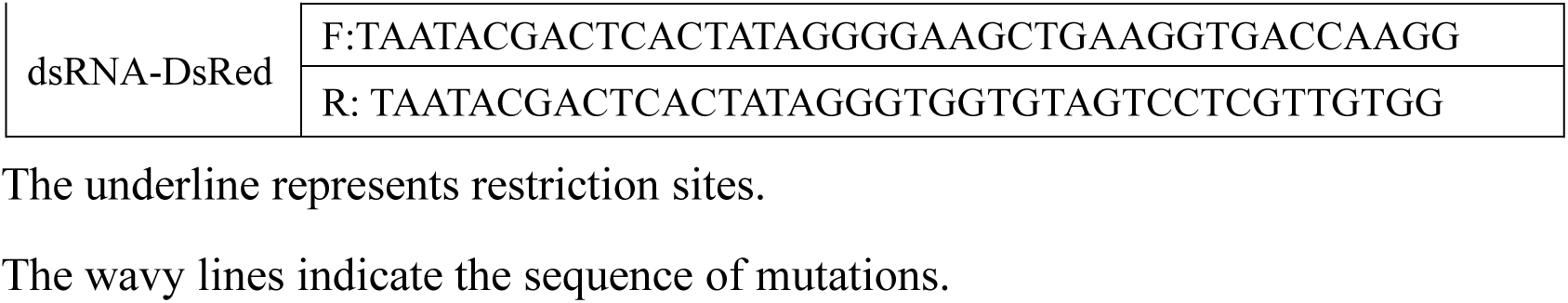
Primers used in the study.

## Notes

### Competing Interest Statement

The authors have declared no competing interest.

